# Cooperative riboregulation in living cells through allosteric programming of toehold activation

**DOI:** 10.1101/009688

**Authors:** Guillermo Rodrigo, Satya Prakash, Shensi Shen, Eszter Majer, José-Antonio Daròs, Alfonso Jaramillo

## Abstract

Living cells rely on small non-coding RNAs (sRNAs) to regulate gene expression at the post-transcriptional level. Contrary to most protein-based activators of transcription, all known riboregulators do not exploit cooperative binding mechanisms to activate gene expression. We conceived a general strategy to design cooperative riboregulation by programming a hierarchical toehold activation cascade, which we implemented into a *de novo* sequence design algorithm. We engineered different riboregulatory systems relying on the conditional formation of a heterotrimeric species. We characterized the specificity of each RNA-RNA interaction *in vitro* and the cooperative activation of gene expression in *Escherichia coli*. As we only rely on a biochemical model to compute allosteric regulation, our strategy could be applied to reach more complex RNA-based nanostructures regulating gene expression for synthetic biology applications.

## Introduction

Living systems rely on cooperative interactions at the molecular level to sustain complex behaviors [1]. Indeed, the control mechanisms of gene expression found in higher organisms (e.g., mammals) present many cellular factors at play and are highly combinatorial [2], but mostly limited to protein interactions. As nucleic acids are molecules with more programmable interactions than proteins, nanotechnology has relied on them to engineer complex nanostructures, conceived by designing interactions using toehold-mediated strand-displacement reactions [3, 4]. However, RNAs have not yet been engineered to control gene expression in a sophisticated manner in living cells.

In this work, we considered RNA as a substrate to design and characterize cooperative riboregulators of gene expression, exploiting a physicochemical model involving free energies and secondary structures. Previous work on *de novo* design of synthetic regulatory RNAs has allowed the engineering of new systems to control protein expression in living cells, such as those based on riboregulators [5 - 10], riboswitches [11 - 13], and ribozymes [14 - 16]. Here, we went beyond by designing obligate heterodimers of small RNAs (sRNAs) activating the initiation of translation. There have not been reported examples of natural or synthetic riboregulators able to regulating gene expression in microbes cooperatively, although this constitutes a road to increase the required nonlinearity to obtain complex behaviors with RNA. Only some examples have been found in higher organisms, mainly regarding the synergistic repression of microRNAs with target sites optimally separated (between 8 and 40 nucleotides) [17]. Our work, in addition to illustrating the designability of artificial cooperative sRNAs, provides experimental evidence to look for combinatorial mechanisms within the microbial riboregulome.

The engineering of cooperative riboregulators is essential to increase the nonlinearity of an RNA system. And this is important because many different useful functions emerge from nonlinear interactions in regulatory circuits. This is case, for instance, of toggle switches [18] or oscillators [19] (based on transcriptional control). However, sRNAs work as monomers, which prevents at first sight its use as a substrate to engineer complex behaviors (e.g., an approximate linear trend of activation resulted between the natural riboregulator DsrA and its target RpoS [20]). The combination of different riboregulations (also applicable for monomeric transcription factors) can nevertheless surmount this issue. We know that nonlinearity can increase through regulatory cascades [21], but this does not occur in case of monomers (Fig. 1a; see also Materials and Methods). Hence, to have a nonlinear dose-response curve with sRNAs (like the one produced by transcription factors that dimerize), we would need the formation of heterodimers for synergistic activation (Fig 1b; see for instance ref. [22] for the case of proteins).

**Figure 1.**
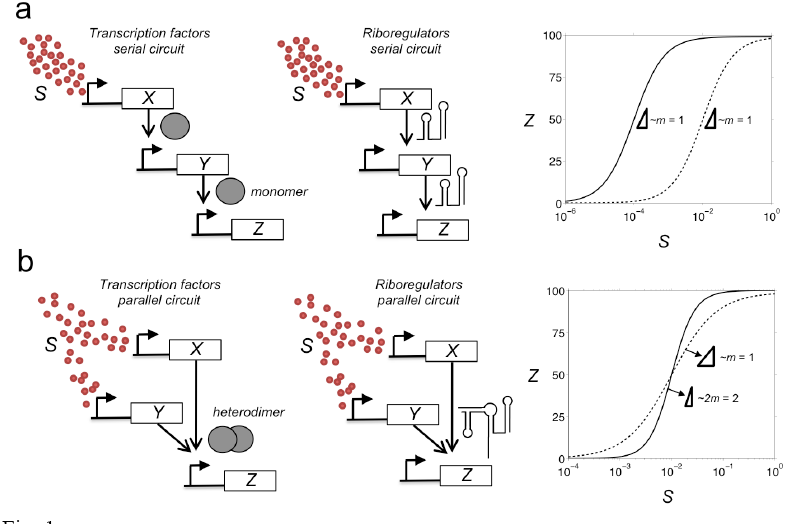
Schemes and model simulations of regulatory cascades of activation. (a) Scenario of serial activation by monomeric transcription factors or riboregulators together with the prediction of the dose-response curve. (b) Scenario of parallel activation by heterodimeric transcription factors or riboregulators together with the prediction of the dose-response curve. In both cases, solid line represents the expression of gene *Z*, whilst dashed line corresponds to a reduced system without gene *Y*. Here we considered *A* = 100, *m* = 1, and *n* = 1 (see Materials and Methods).

## Results and Discussion

To design cooperative riboregulation, we exploited the fact that *i*) RNA-RNA interactions are kinetically dominated by a toehold-mediated mechanism (i.e., interacting regions unpaired within the intramolecular structures of the species), *ii*) the ribosome-binding site (RBS) of a given mRNA can be considered as a toehold mediating the interaction with the ribosome (this is exploited in many natural/synthetic systems [5, 12, 14]), and *iii*) activating toeholds in cascades implement a hierarchical assembly of higher-order complexes [23]. We propose programming RNA species holding hidden/inactive toehold domains that become active after a conformational change following a binary interaction. Once the toehold is active, the complex is able to interact with a third RNA species. The energy landscape associated to our hypothesized hierarchical interaction mechanism is shown in Figure 2 (see also Fig. S3).

**Figure 2.**
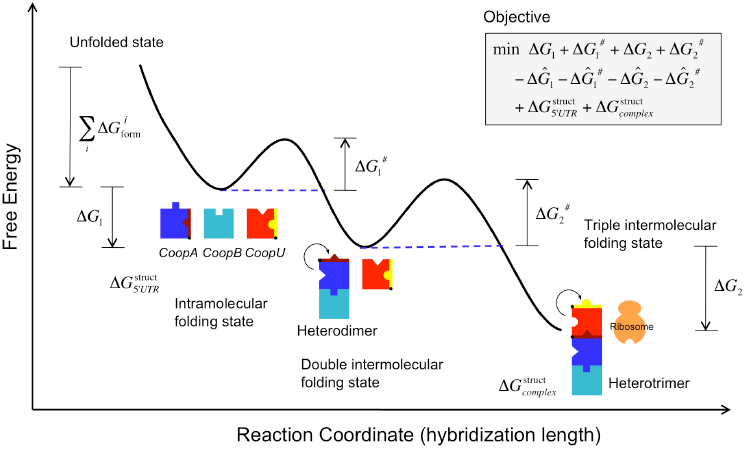
Energy landscape of cooperative riboregulation of gene expression in bacterial cells. On left top, the objective function to be minimized is explicited. The terms Δ*G*_1_, 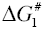, Δ*G*_2_, and 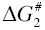 correspond to the free energies of hybridization and activation, respectively, between the two sRNAs (subscript 1) and between the sRNA complex (heterodimer) and the 5’ UTR (subscript 2). Note that the free energy of hybridization is a negative magnitude, whereas the free energy of activation is a positive magnitude. The terms 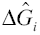 and 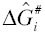 correspond to the free energies of hybridization and activation, respectively, between the sRNA *i* and the 5’ UTR (subscript *i* = 1 or 2, for CoopA or CoopB). They have to be maximized, so they have a minus sign. Finally, the terms 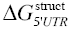 and 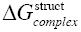 are the works required to fold the 5’ UTR (CoopU) and the heterotrimer according to the structure specifications (in this case, RBS paired in the 5’ UTR and unpaired in the complex). On right top, the scheme of the optimization loop, where three RNA sequences (CoopA, CoopB, and CoopU) are iteratively mutated and evaluated according to the objective function, is shown. The energy landscape shows the different conformational states (intra-and intermolecular), together with the free energy terms of the objective function, as a function of a reaction coordinate (number of intermolecular base pairs). The trajectory illustrates the interaction between two sRNAs to form a heterodimer that is able to subsequently interact with the 5’ UTR to release the RBS and then allow translation.

To control gene expression, we used *trans*-acting sRNAs able to activate a *cis*-repressed 5’ untranslated region (UTR) of a given messenger RNA (mRNA), with a mechanism of allosteric regulation [5]. We identified the different conformational states and their free energy levels, which could be predicted with a physicochemical RNA model [24 - 27]. The reaction coordinate was defined as the number of intermolecular hydrogen bonds (or base pairs), on one side, between the two sRNAs (CoopA and CoopB hereafter) and, on the other side, between the resulting sRNA complex and the 5’ UTR of the target mRNA (CoopU hereafter). On the energy landscape, two barriers (free energies of activation; 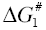 and 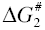) impinge the progression of the reaction, one for each intermolecular interaction that defines the reaction coordinate. These free energies of activation are associated to the degree of exposition of the toeholds to the solvent, and have to be low (i.e., three or more nucleotides unpaired in the toehold [23]) to permit the initiation of the reaction (kinetic aspect). In addition, for an efficient reaction, the free energies of hybridization (Δ*G*_1_ and Δ*G*_2_) have to be as low as possible (at least, lower than -15 Kcal/mol [23]) to ensure irreversibility in the intermolecular interactions (thermodynamic aspect) [28].

To show that the proposed hierarchical toehold activation mechanism (based on the kinetic aspect) is a sufficient criterion to provide arbitrary cooperative riboregulation, we developed a computational algorithm that addressed the *de novo* sequence design (iterative process of random mutations and selection according to an energy-based objective function) (Fig. S2; see also Materials and Methods) [9, 23]. The objective function was calculated with a nucleotide-level energy model considering all conformational states of the system’s species (CoopA, CoopB, CoopU, all possible heterodimers, and the heterotrimer), following a combined strategy of positive and negative design [29]. On the one hand, as positive objectives (to be minimized), we considered the free energies of activation and hybridization corresponding to the interactions between the two sRNAs and between the resulting sRNA complex and the 5’ UTR. We also considered the intramolecular structure of the 5’ UTR (to have the RBS paired), and the intermolecular structure of the 5’ UTR with the sRNA complex (to have the RBS unpaired). On the other hand, as negative objectives (to be maximized), we took the free energies of activation and hybridization corresponding to the interactions between each sRNA and the 5’ UTR.

We designed computationally two systems: Coop1 and Coop2 (Table S1). System Coop1 was obtained by specifying the intramolecular structures of the two sRNAs, whereas for system Coop2 the specification was of the intermolecular structure of the sRNA complex (introduced as soft constraints in both cases). These specifications, although not functionally required, were introduced to prevent premature degradation of unstructured sRNAs. For these systems, the toehold that nucleates the interaction between CoopA1:CoopB1 and CoopU1 (or CoopA2:CoopB2 and CoopU2) is hidden within the intramolecular structure of CoopA1 (or CoopA2). However, the toehold that nucleates the interaction between CoopA1 and CoopB1 (or CoopA2 and CoopB2) is unpaired (active) within their intramolecular structures, so the reaction of hybridization between both sRNAs can take place. As a result, within the intermolecular structure of the resulting sRNA complex, the toehold that nucleates the interaction with CoopU1 (or CoopU2) becomes active. The structures of system Coop2 are depicted in Figure 3a, showing the hierarchical activation of toeholds.

**Figure 3.**
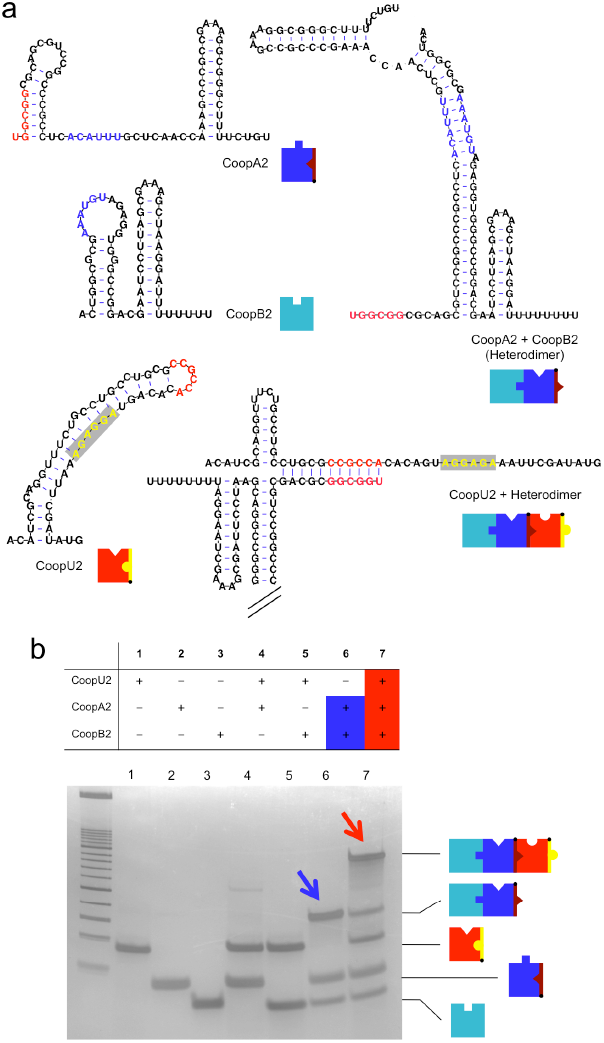
Molecular characterization of cooperative riboregulators *in vitro*. (a) Sequences and structures of the species of the designed sRNA system Coop2. The toehold for the interaction between the two sRNAs is shown in blue. The toehold for the interaction between the heterodimer (sRNA complex) and the 5’ UTR is shown in red. In the 5’ UTR (CoopU2), the RBS is shown in yellow. The transcription terminators T500 and B1002 were used in CoopA2 and CoopB2, respectively. (b) Electrophoretic analysis showing the hierarchical interaction between sRNAs. The formation of the heterodimer and the heterotrimer are marked by arrows.

However, even if a toehold is not hidden within the corresponding intramolecular structure (of CoopA), it may still remain inactive. In this case, the hybridization free energy would not be sufficient to ensure irreversible interaction (with CoopU), and an additional species (CoopB) would be required for the reaction. The free energy of hybridization between CoopA:CoopB and CoopU would then be sufficient to form the triple intermolecular folding state with a three-way junction. To explore this possibility (based on the thermodynamic aspect), we considered a design based on a three-way junction; indeed natural occurring three-way junctions have been already exploited to engineer RNA-based structures [30], although not to control gene expression. We took advantage of our previously published riboregulatory system RAJ11 [9]. We split the sRNA into two halves, CoopA11 and CoopB11 hereafter, and considered the cognate 5’ UTR, which we named here CoopU11 (Fig. S4). The free energies of hybridization were automatically fair thanks to the three-way junction formed within the intermolecular conformation between the sRNA and 5’ UTR in the native system RAJ11. When constructing CoopA11 and CoopB11, we found that both sRNAs had an active toehold that allowed them to interact. The heterodimer CoopA11:CoopB11 has another active toehold that nucleates its binding to CoopU11 by forming a heterotrimer with the three-way junction. This resulting structure activates the RBS for recognition by the ribosome and then initiation of translation.

We performed in a PAGE gel the molecular characterization of the higher-order RNA-RNA interactions [30]. The complementary DNAs corresponding to the RNA species were first transcribed *in vitro* (for the sRNA species without transcription terminators). We prepared in each lane the three individual species: CoopU, CoopA, and CoopB. Then, we prepared the three possible combinations of two species, and, finally, the three species together. The gel finely revealed, for system Coop2, the intermolecular interactions *i*) between the two sRNAs, and *ii*) between the resulting sRNA complex and the 5’ UTR (Fig. 2b, lanes 6 and 7). It also revealed mild intermolecular interaction between one sRNA and the 5’ UTR (Fig. 3b, lane 4). We also confirmed in a PAGE gel the intermolecular interaction between the sRNA complex and the 5’ UTR for system Coop11 (Fig. S5). Taken together, these results validated our description of the energy landscape.

The designed RNA systems were implemented as separate operons (with their respective transcriptional terminators) in plasmids (Fig. S1), which were transformed into *E. coli* cells (see Materials and Methods) expressing the transcriptional repressors LacI and TetR (Fig. 4a illustrates the engineered sRNA circuit). The use of P_L_-based inducible promoters allowed controlling the expression of the two sRNAs with the external inducers isopropyl-β-D-thiogalactopyranoside (IPTG) and anhydrotetracycline (aTc) [31]. Figure 4b shows the dynamic ranges (characterized by fluorometry) of our engineered systems, probing the regulation of gene expression in living cells with two cooperative sRNAs (at the population level). Table S3 shows eventual off-target effects of our designed riboregulators using RNApredator [32] (considering the 5’ UTRs of all mRNAs in the genome of *E. coli* K-12 MG1655), although the viability of the living cell is not compromised when expressing our systems. To assess that the combined action of the two sRNAs was enhanced over the expected action from their independent contributions, we compared the increase in gene expression with both inducers with respect to the additive increase with IPTG, on one side, and aTc, on the other (Welch *t*-test, *P* < 0.02 for systems Coop11 and Coop2; and *P* = 0.06 for system Coop1). We also observed that *cis*-repression is much stronger in systems Coop11 and Coop1 than in system Coop2 (Fig. S6). Figure 4c demonstrated a graded response with both inducers. To further explore the cooperative behavior at single cell level, we performed a time-dependent characterization of system Coop2 using microfluidics devices (Fig. S7) [33]. This allowed us to monitor GFP expression in single cells under a varying concentration of both inducers (Fig. 4d). These results showed that each individual cell responded to the inducers, and that the system, as expected, is reversible *in vivo*. Flow cytometry experiments also revealed significant population shift in response to both inducers (Figs. 4e and S8).

**Figure 4.**
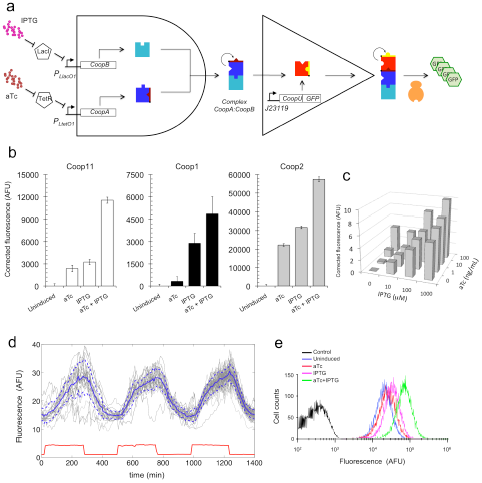
Functional characterization of designed cooperative riboregulatory systems in bacterial cells. (a) Digital-like scheme of the engineered sRNA circuit. Promoters P_LlacO1_ and P_LtetO1_ control the expression of the two sRNAs, which can be tuned with external inducers IPTG and aTc, whereas the mRNA is constitutively expressed from promoter J23119. The two sRNAs first interact to form a complex that is then able to activate a *cis*-repressed gene. The reporter gene is a GFP. (b) Fluorescence results of the designed sRNA systems Coop11, Coop1, and Coop2 for all possible combinations of inducers. Error bars represent standard deviations over replicates. (c) Fluorescence results for system Coop2 with a gradient of IPTG and aTc. (d) Single cell tracking of fluorescence [GFP expression in arbitrary fluorescence units (AFU)] in one microchamber of the microfluidics device under time-dependent induction with IPTG (1 mM) and aTc (100 ng/mL) for system Coop2. A square wave of both inducers with period 8 h (i.e., 4 h induction and 4 h relaxation, ON/OFF) was applied. The solid and dashed lines (in blue) correspond to the mean and plus/minus standard deviation over the cell population. Sulforhodamine B was used to monitor the inducer time-dependent profile (in red). (e) Flow-cytometric results for system Coop2 for all possible combinations of inducers. The black curve corresponds to plain cells (without plasmid; control).

In conclusion, we programmed with RNA a hierarchical activation of hidden toeholds after intermolecular interactions (systems Coop1 and Coop2; sequences obtained by computational design). Cooperative behavior can also be programmed without relying on the activation of a hidden toehold when forming the obligate heterodimer if a thermodynamically stable three-way junction can be used (system Coop11). This coupling between hierarchical assembly of RNA structures with gene expression could allow the application of known nanotechnology structures [34, 35] to engineer higher-order regulation of gene expression in living cells. This will contribute to have building blocks with increased nonlinearity, and potentially to create novel RNA circuits with sophisticated functionalities.

## Materials and Methods

### Modeling simple regulatory cascades of activation

We considered a simple regulatory scenario of a cascade of activation by regulators that can be either transcription factors or riboregulators. A basic Hill-like model was used to account for activation of gene expression [18, 19]. An environmental signal (*S*), working with effective cooperativity *m*, was considered to trigger the cascade, which lead us to write

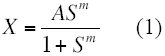

where *X* is the expression of the regulated gene and *A* an integrative parameter for normalization purposes (typically *A* = 10 - 1000). Then, a new gene (*Z*) is activated in a subsequent step, given by

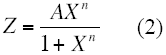

where *n* is the self-cooperativity of the regulator (i.e., *n* = 1 if it does not self-aggregate to form dimers or higher-order elements).

By considering *Y* another regulator at play, we could have two scenarios: one in which *S* activates *X*, which activates *Y*, which then activates *Z* (regulation in series), and another in which *S* activates *X* and *Y*, which then both activate *Z* (regulation in parallel). In the first case, we obtained 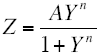 [with *Y* given by Eq. (2) and *X* by Eq. (1)]. Therefore, when *n* = 1, it turned out

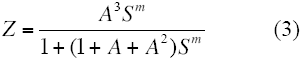

where there is no increase in effective cooperativity between *Z* (output) and *S* (input), neither if we increase the length of the cascade. This only happens when *n* > 1. However, we had 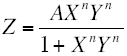 in the second case by considering synergistic activation [with *X* and *Y* given by Eq. (1)], and this gave (again for *n* = 1)

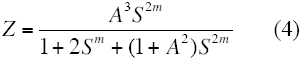

highlighting an increase in effective cooperativity between *Z* and *S* (from *m* to 2*m*).

### RNA sequence design

We developed a Monte Carlo Simulated Annealing optimization algorithm [36] to design regulatory RNAs that cooperate to regulate gene expression. The system was composed of three different RNA species: two small RNAs (sRNAs) and one 5’ untranslated region (UTR). To implement this algorithm, we constructed a physicochemical model based on free energies and RNA structures [25] that involved, amongst, the energies of activation and hybridization corresponding to the interaction between the two sRNAs and the energies of activation and hybridization corresponding to the interaction between the sRNA complex and the 5’ UTR. The model also accounted for the degree of repression and exposition of the RBS within the 5’ UTR intramolecular and intermolecular structures. Rounds of random mutations were applied and selected with the energy-based objective function (Fig. S2). For that, we extended a previously reported algorithm for RNA design [9]. We used the Vienna RNA package [24] for energy and structure calculation. The sequences engineered in this work of cooperative riboregulators, as well as their cognate 5’ UTRs, are shown in Table S1.

### Plasmid construction

The different sRNA systems were chemically synthesized (IDT) and cloned in a pSTC2-based plasmid that contained a pSC101m replication origin (a mutated pSC101 ori giving a high copy number) and a kanamycin resistance marker (Fig. S1). The pSTC2 vector, used in our previous works, has a superfolder Green Fluorescent Protein (sfGFP) [37] as reporter gene, with a *ssr*A degradation tag [38] for a fast turnover. The promoters P_LlacO1_ and P_LtetO1_ [31] control the expression of the two sRNAs, whereas the messenger RNA (containing the 5’ UTR) is constitutively expressed from promoter J23119. Strains and plasmids used in this study are listed in Table S2.

### Cell culture and reagents

*Escherichia coli* strain DH5α (Invitrogen) was used for plasmid construction purposes as described in the manual [39]. Characterization experiments were performed in *E. coli* DH5α-Z1 cells (Clontech) or in *E. coli* K-12 MG1655-Z1 cells (both *lacI*^+^ *tetR*^+^) for control over the promoters P_LlacO1_ and P_LtetO1_. Plasmids carrying systems Coop11, Coop1 and Coop2 were transformed into DH5α-Z1 cells for characterization in fluorometer (TECAN). Plasmid carrying system Coop2 was transformed into MG1655-Z1 cells for characterization in a microfluidic device, and into DH5α-Z1 cells for characterization in a flow cytometer.

Cells were grown aerobically in LB medium or in M9 minimal medium, prepared with M9 salts (Sigma-Aldrich), glycerol (0.8%, vol/vol) as only carbon source, CaCl_2_ (100 μM), MgSO_4_ (2 mM), and FeSO_4_ (100 μM). Kanamycin concentration was 50 μg/mL. Cultures were grown overnight at 37 °C and at 225 rpm from single-colony isolates before being diluted for *in vivo* characterization. 1 mM IPTG (Thermo Scientific) was used for full activation of promoter P_LlacO1_ when needed, and 100 ng/mL aTc (Sigma-Aldrich) was used for full activation of promoter P_LtetO1_. For microfluidics cell cultures, cells were grown aerobically in fresh LB and in LB supplemented with 0.05% sulforhodamine B (Sigma-Aldrich), and IPTG + aTc (i.e., we used sulforhodamine B to monitor the presence of inducers in the chamber).

### *In vitro* RNA interaction

We first constructed the cDNAs of the different RNA species of the designed system to then perform the *in vitro* transcription. We analyzed the systems Coop2 and Coop11. We considered the sRNAs without transcription terminators and the 5’ UTR until the start codon. Amplification by PCR (30 cycles, extension 0:30 min), using Phusion DNA polymerase (Thermo Scientific), was done over the template plasmid (pCoop2 or pCoop11). The PCR products were cloned into the plasmid pUC18, where the restriction site Eco31I was previously removed. The resulting plasmids with inserts were selected by DNA cleavage with appropriate restriction enzymes. Sequences were also verified by sequencing.

To perform the *in vitro* transcription, 3 μg of each plasmid was digested with Eco31I (4 μL, own preparation in the lab), and purified with silica-based columns (Zymo). We used approximately 1 μg of digested plasmid in the reaction. This was in 20 μL: 10 μL of plasmid, 2 μL Buffer 10x (Roche), 0.4 μL DTT 10 mM, 1 μL NTPs 10 mM (Fermentas), 0.5 μL Ribolock (∼40 U/μL, Thermo Scientific), 1 μL PPase (0.1 U/μL, Fermentas), 1 μL RNA polyerase T7 (50 U/μL, Epicentre), and 4.1 μL H2O. We incubated the mix for 1 h at 37 °C, and then added 20 μL of it with formamide. The samples were heated at 95 °C for 1:30 min, then cooled on ice, and then loaded in a gel PAGE 10% 8 M urea, TBE (1x), (200 V, 2:30 h). We cut the resulting bands for purification. We confirmed the presence of RNA by loading a small part of purifications in another gel PAGE 10% 8 M urea, TBE (1x) (200V, 2:30 h).

For the reaction of RNA-RNA interaction, we used approximately 20 ng of RNA for each of the trascripts. The buffer of the reaction was: 50 mM Tris-HCl pH 7.5, 10 mM MgCl2, 20 mM NaCl. The reactions (20 μL) were denaturalized (1.5 min at 95 °C) and slowly cooled (15 min at room temperature) [30]. We then added 1.5 μL glycerol (87%) and 0.2 μL AB-XC (100x) to load the gel (15% PAGE, TAE, 1 mm), which was run for 2 h with 75 mA at 4 °C. The gel was dyed with ethidium bromide and silver (see e.g. Fig. S5). We used the DNA molecular weight marker XIII (50 bp ladder, Roche).

### Fluorescence quantification

Cells were grown overnight in LB medium, and were then refreshed by diluting 1:200 in M9 medium. They were grown for additional 2 h to then load 200 μL in each well of the plate (Custom Corning Costar). Appropriate inducers (none, aTc, IPTG, or aTc + IPTG) were introduced when needed. The plate was incubated in an Infinite F500 multi-well fluorometer (TECAN) at 37 °C with shaking. It was assayed with an automatic repeating protocol of absorbance measurements (600 nm absorbance filter) and fluorescence measurements (480/20 nm excitation filter - 530/25 nm emission filter for sfGFP) every 15 min. All samples were present in triplicate on the plate.

Normalized fluorescence was obtained by subtracting the background values corresponding to M9 medium (in both fluorescence and absorbance values) and then dividing fluorescence by absorbance at OD_600_ ≈ 0.5. Corrected fluorescence was obtained by subtracting the fluorescence of uninduced cells.

### Single cell microfluidic analysis

The design of our microfluidic device was performed in AUTOCAD (AUTODESK) and was already applied to study a synthetic genetic oscillator [40]. All images were acquired using Zeiss Axio Observer Z1 microscopy (Zeiss). The microscope resolution was 0.24 μm with Optovariation 1.6X, resulting total magnification 1600X for both bright field and fluorescent images. Images were analyzed with MATLAB (MathWorks). Cells were tracked by defining a cell-to-cell distance matrix and the cell lineages were reconstructed. Finally, the fluorescence level of each cell in each fluorescence frame was extracted (see Fig. S7 for the setup).

### Flow cytometry analysis

Cells were grown overnight in LB medium, and were then diluted 1:200 in fresh LB medium containing inducers (none, aTc, IPTG, or aTc + IPTG) and incubated to reach an OD_600_ of 0.2-0.4. Afterwards, cells were diluted again in 1 mL PBS. All expression data were analyzed using a Becton-Dickinson FACScan flow cytometer with a 488 nm argon laser for excitation and a 530/30 nm emission filter (GFP). Gene expression of each sample was obtained by measuring the fluorescence intensity of thousands of cells. Data were analyzed using the FlowJo software by gating the events using scatter ranges (see Fig. 8). Fluorescence distribution is presented in log scale.

## Supplementary Information

Supplementary information is available.

## Acknowledgements

Work supported by grants FP7-ICT-610730 (EVOPROG) and FP7-KBBE-613745 (PROMYS) to A.J., and BIO2011-26741 (Ministerio de Economía y Competitividad, Spain) to J.-A.D. G.R. is supported by the AXA Research Fund, and E.M. by a predoctoral fellowship (AP2012-3751, Ministerio de Educación, Cultura y Deporte, Spain).

## Author contributions

G.R. and A.J. conceived this study. G.R., S.S., J.-A.D., and A.J. designed experiments. G.R., S.P., S.S., and E.M. performed experiments. G.R., S.S., J.-A.D., and A.J. analyzed the results. G.R. and A.J. wrote the manuscript.

## Supplementary Information

### Supplementary Figures

**Figure S1:**
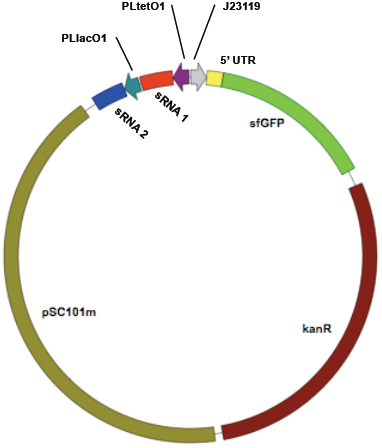
Map of the plasmid used in this work for expressing the designed sRNA systems. The sRNAs 1 and 2 correspond to CoopA and CoopB, respectively, according to our terminology. The 5’ UTR is named as CoopU.

**Figure S2:**
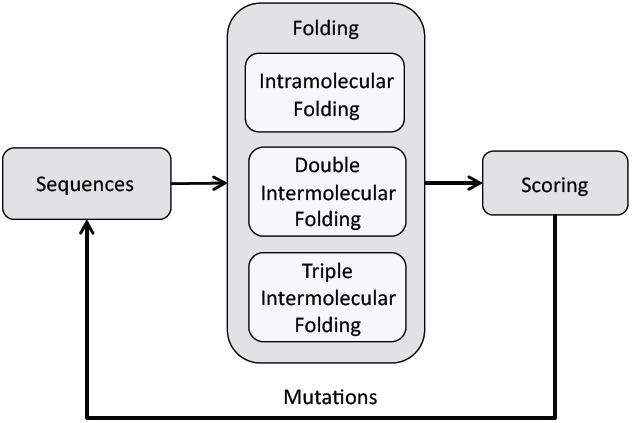
Scheme of the optimization loop, where three RNA sequences (CoopA, CoopB, and CoopU) are iteratively mutated and evaluated according to the objective function. To fold the sequences we used ViennaRNA [1].

**Figure S3:**
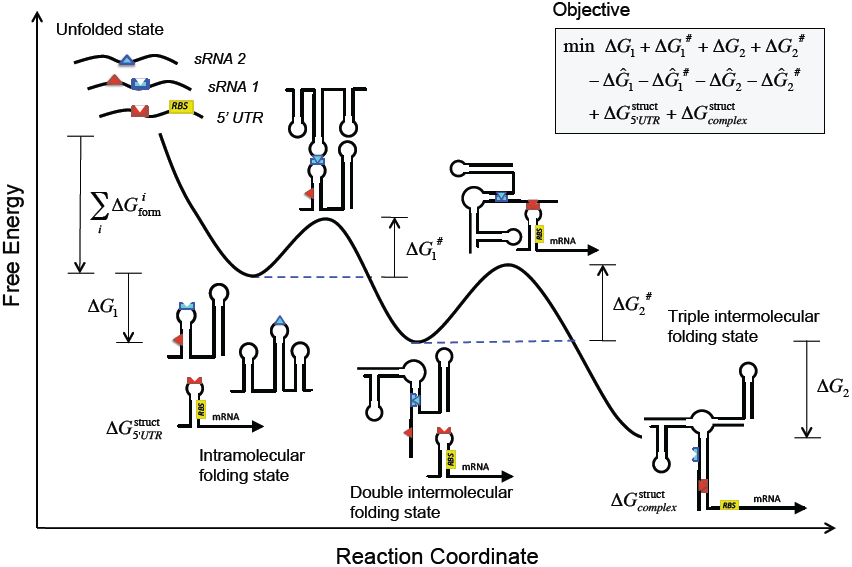
Energy landscape of cooperative riboregulation of gene expression in bacterial cells. On left top, the objective function to be minimized is explicited. The terms Δ*G*_1_, 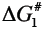, Δ*G*_2_, and 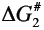 correspond to the free energies of hybridization and activation, respectively, between the two sRNAs (subscript 1) and between the sRNA complex and the 5’ UTR (subscript 2). Note that the free energy of hybridization is a negative magnitude, whereas the free energy of activation is a positive magnitude. The terms 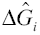 and 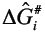 correspond to the free energies of hybridization and activation, respectively, between the sRNA *i* and the 5’ UTR (subscript *i* = 1 or 2). They have to be maximized, so they have a minus sign. Finally, the terms 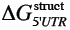 and 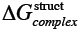 are the works required to fold the 5’ UTR and the complex (three RNAs) according to the structure specifications (in this case, RBS paired in the 5’ UTR and unpaired in the complex). The energy landscape shows the different conformational states (intra- and intermolecular), together with the free energy terms of the objective function, as a function of a reaction coordinate (number of intermolecular base pairs). The trajectory illustrates the interaction between two sRNAs to form a sRNA complex that is able to subsequently interact with the 5’ UTR to release the RBS and then allow translation.

**Figure S4:**
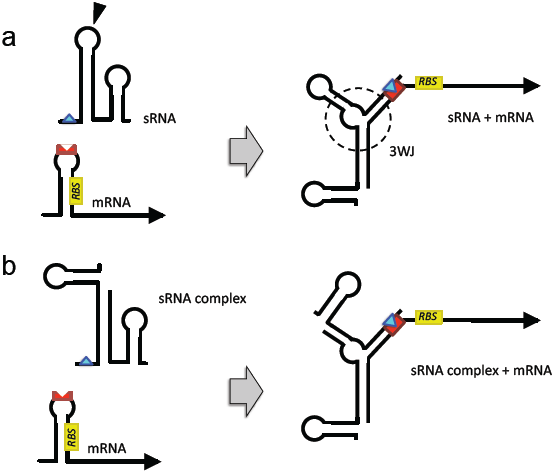
(a) Scheme of the riboregulatory system RAJ11 (one sRNA interacts with the 5’ UTR of mRNA) [2]. (b) Scheme of the cooperative riboregulatory system Coop11 (two sRNAs form a complex that interacts with the 5’ UTR). This system is based on the previous one by taking advantage of the three-way junction (3WJ) formed to then split the sRNA in two at the wedge (and add a terminator to the first fragment). The sRNAs are illustrated with terminators.

**Figure S5:**
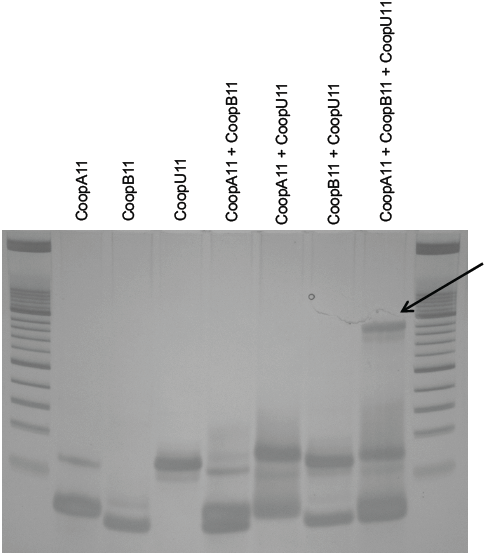
Electrophoretic analysis of system Coop11. The different lanes correspond to all combinations of species. The arrow marks the interaction of the three RNAs.

**Figure S6:**
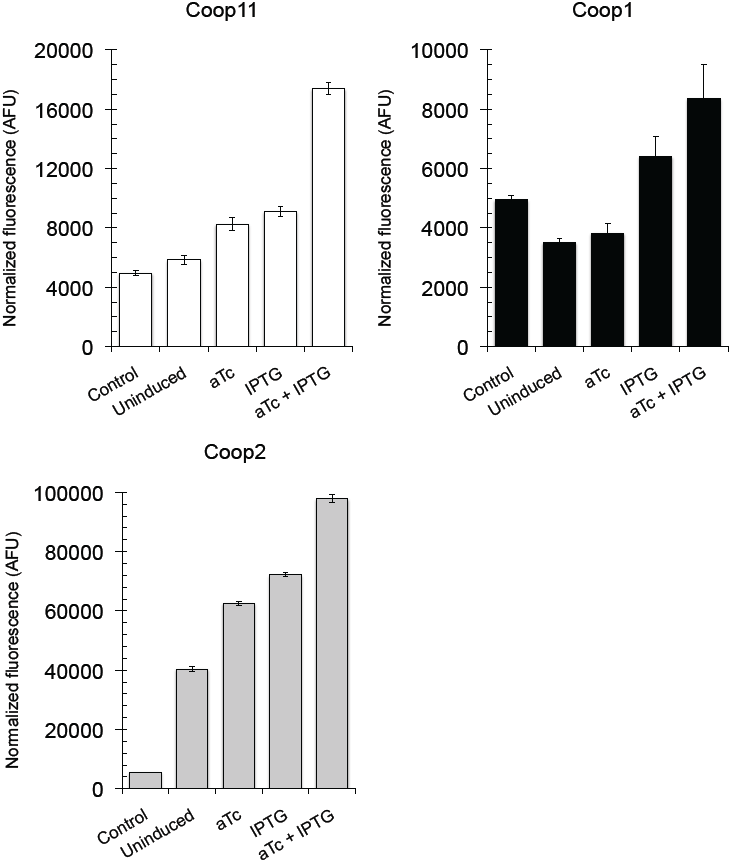
Fluorescence results of the designed sRNA systems Coop11, Coop1, and Coop2 for all possible combinations of inducers. Error bars represent standard deviations over replicates. Uninduced corresponds to cells transformed with the plasmid but without inducers in the medium, whilst control corresponds to plain cells without plasmid. These results show that system Coop2 has a higher basal translation rate (i.e., *cis*-repression is much stronger in systems Coop11 and Coop1 than in system Coop2).

**Figure S7:**
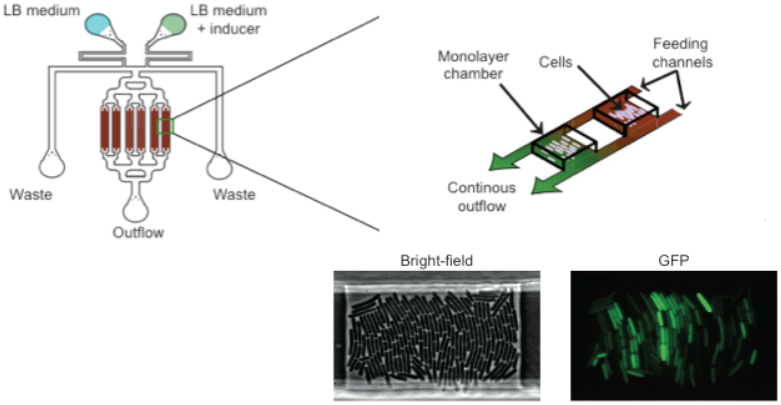
Scheme of the microfluidic device used to monitor GFP expression in single cells.

**Figure S8:**
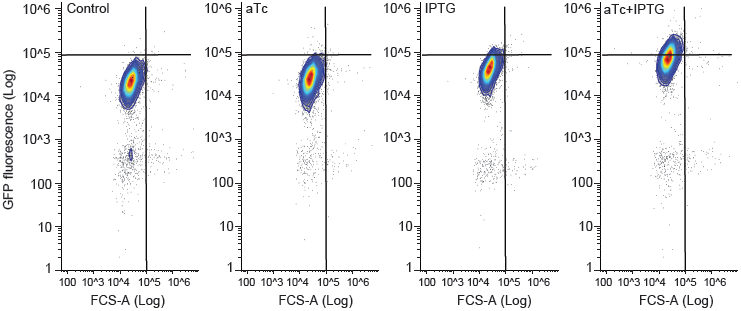
Scatter plots of flow cytometry for system Coop2.

### Supplementary Tabels

**Tabel S1:**
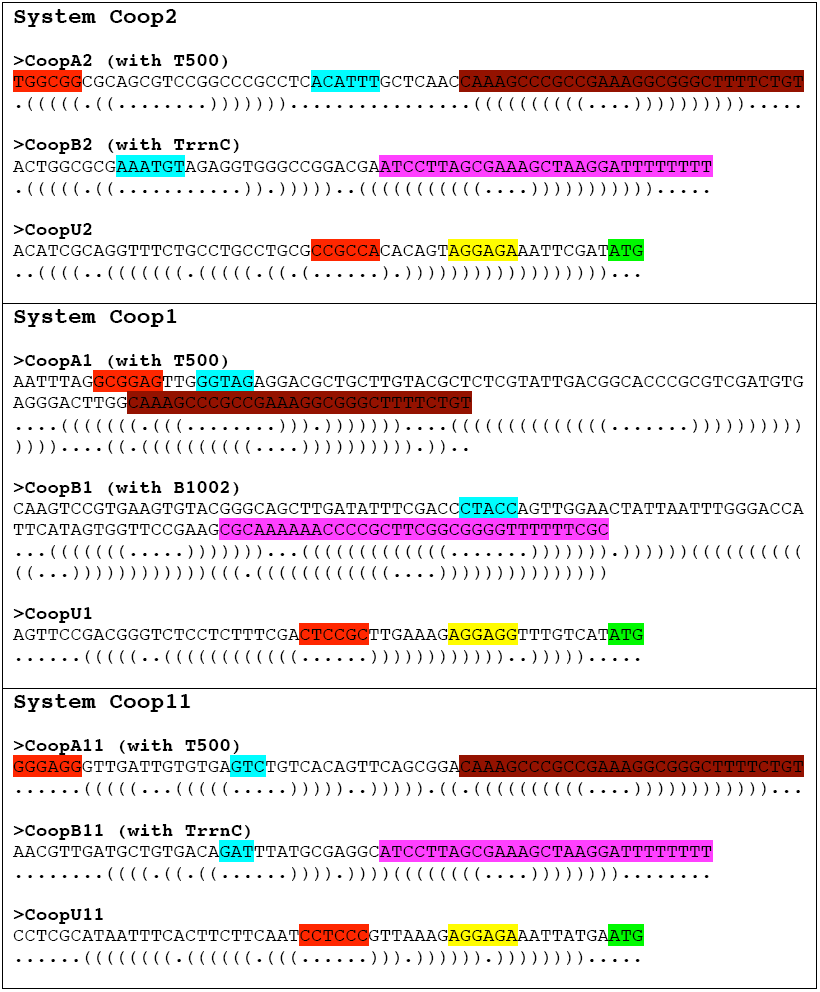
Sequences of the cooperative sRNA systems designed in this work. Dot-bracket structures are also shown. The seed region for the interaction between the two sRNAs (CoopA and CoopB) is shown in cyan. The seed region for the interaction between the sRNA complex and the 5’ UTR (CoopU) is shown in red. In CoopU, the RBS is shown in yellow and the start codon in green. The transcription terminator T500 (efficiency > 90%) was used in CoopA, shown in dark red, and the terminator TrrnC (efficiency > 90%) or B1002 (efficiency about 90%) in CoopB, shown in magenta.

**Table S2:** Strains and plasmids used in this work.

| Strains or plasmids | Features | Ref. |
| --- | --- | --- |
| *E. coli* DH5α | Commercial | Invitrogen |
| *E. coli* DH5α-Z1 | Commercial (DH5α, *lacIQ*, PN25-*tetR*, SpR) | Clontech |
| *E. coli* MG1655-Z1 | *lacIQ*, PN25-*tetR*, SpR | Gifted by<br>M.B. Elowitz |
| pCoop2 | pSC101m ori, kanR, sfGFP-LAA | This work |
| pCoop1 | pSC101m ori, kanR, sfGFP-LAA | This work |
| pCoop11 | pSC101m ori, kanR, sfGFP-LAA | This work |
| pRAJ11 | pUC ori, ampR-kanR, GFPmut3b | [2] |

**Table S3:**
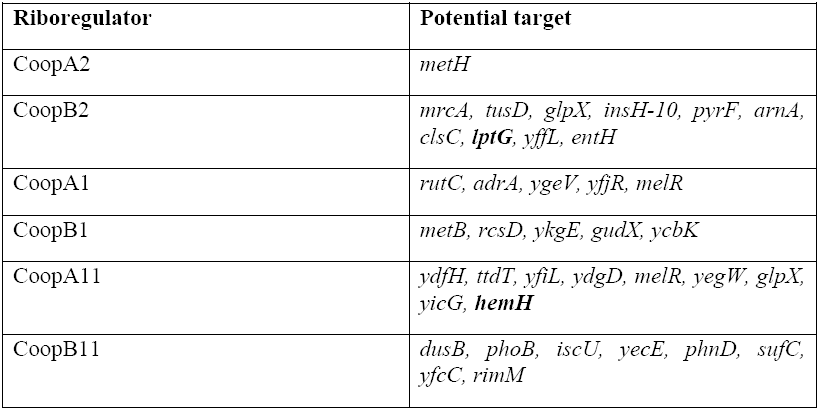
Prediction of eventual off-target effect of the designed sRNAs using RNApredator [3]. Neighborhood of 90 nt before and 10 nt after the start codon (in *E. coli* K-12 MG1655). Essential genes bold-faced [4] (although the sRNAs do not hybridize with the RBSs of the essential genes targeted).

